# Structural Features and Oligomeric Nature of Human Podocin Domain

**DOI:** 10.1101/2020.05.21.108415

**Authors:** Sandeep KN Mulukala, Shivkumar S Irukuvajjula, Krishan Kumar, Kanchan Garai, Pannuru Venkatesu, Ramakrishna Vadrevu, Anil K Pasupulati

**Affiliations:** Department of Biochemistry, School of Life Sciences, University of Hyderabad, Hyderabad, India – 500046; Department of Biological Sciences, Birla Institute of Technology and Sciences- Pilani Hyderabad Campus, Hyderabad, India – 500078; Department of Chemistry, University of Delhi, New Delhi, India - 110 007; Tata Institute of Fundamental Research, Hyderabad, India - 500019

**Keywords:** Podocin, Podocyte, Nephrotic syndrome, slit-diaphragm, proteinuria

## Abstract

Podocytes are crucial cells of the glomerular filtration unit and playing a vital role at the interface of the blood-urine barrier. Podocyte slit-diaphragm is a modified tight junction that facilitates size and shape-dependent permselectivity. Several proteins including podocin, nephrin, CD2AP, and TRPC6 form a macromolecular assembly and constitute the slit-diaphragm. Podocin is an integral membrane protein attached to the inner membrane of the podocyte via a short transmembrane region (101-125). The cytosolic N- and C-terminus help podocin to attain a hook-like structure. Podocin shares 44% homology with stomatin family proteins and similar to the stomatin proteins, podocin was shown to associate into higher-order oligomers at the site of slit-diaphragm. However, the stoichiometry of the homo-oligomers and how it partakes in the macromolecular assemblies with other slit-diaphragm proteins remains elusive. Here we investigated the oligomeric propensity of a truncated podocin construct (residues:126-350). We show that the podocin domain majorly homo-oligomerize into a 16mer. Circular dichroism and fluorescence spectroscopy suggest that the 16mer oligomer has considerable secondary structure and moderate tertiary packing.

## Introduction

Vertebrate kidneys regulate electrolyte and water balance to maintain body homeostasis. Each kidney is composed of about one million nephrons. The glomerulus and the renal tubule are the two major parts of a nephron that work in unison to ensure protein-free ultra-filtrated urine. The glomerular filtration barrier (GFB) offers permselectivity for the filtration of plasma components into the urine. The GFB consists of fenestrated capillary endothelium, glomerular basement membrane, and podocytes [1]. Podocytes are highly differentiated visceral epithelial cells that encase the glomerular capillaries. A typical podocyte cell consists of a protuberant cell body with primary processes made of actin and microtubules. The primary process further branches into secondary foot processes which interlaces with the neighboring foot processes forming a modified tight junction called the slit-diaphragm (SD) [2]. The SD is a negatively charged zipper-like structure bridging the 30-40nm gap between the adjacent foot processes. This structure curbs the passage of albumin and other large molecules from the blood into primary urine thereby tightly regulating the composition of the glomerular filtrate [3, 4]. The intricate structure of the SD is maintained by an array of protein assemblies. Proteins such as podocin, nephrin, CD-2 associated protein (CD2AP), transient receptor potential cation channel subfamily C-member 6 (TRPC6), zonula occludens-1 (ZO-1), and Nephrin-like proteins 1, 2, and 3 (NEPH) interact to form macromolecular complexes and constitute the structure of SD [5–8].

Mutations in the SD proteins are associated with nephrotic syndrome (NS), which is presented with massive proteinuria, hypoalbuminemia, and edema [9]. Corticoid therapy is the usual recourse to abate NS and based on the patient’s response to corticoid therapy NS is divided into steroid-sensitive NS (SSNS) and steroid-resistant NS (SRNS). Patients with congenital nephropathies usually belong to the SRNS group since they do not respond to corticoid therapy. Congenital nephropathy typically onsets in infants during 0-3 months of age which eventually progresses to irreversible kidney failure within a decade. Mutations in *NPHS1* and *NPHS2* that encode for nephrin and podocin respectively result in the majority of congenital SRNS cases [10]. About 18% of the reported SRNS cases are due to mutations in podocin [11, 12]. Mutations in other SD proteins such as TRPC6 and CD2AP have also been observed to cause NS but at a less frequent rate than nephrin and podocin [13–15].

Podocin is a 383 amino acid protein localizing to the lipid rafts along with other SD proteins [7, 16, 17]. Podocin shares 44% homology with stomatin family proteins due to the presence of a highly conserved PHB domain [18]. Indeed, podocin was predicted to share several structural similarities with the stomatin proteins [5, 18]. Podocin adapts a hook-like structure with its cytoplasmic N- and C-terminus as it attaches to the inner side of the plasma membrane via a transmembrane domain at 100-125 residues [5, 12, 19]. Structural characterization of stomatin revealed that different truncations of the protein associated with different oligomeric states and that the C-terminus of the protein is crucial for homo-oligomerization [20–22]. Studies with truncated C-terminal human podocin revealed that it forms dimers [23]. Though it also proposed that longer constructs of podocin were capable of associating into further higher-order oligomers, it was never demonstrated [23]. Additionally, co-immunoprecipitation studies with nephrin, CD2AP, TRPC6, and NEPH1 indicated that these proteins interact majorly with the C-terminus of podocin [6–9, 16, 17, 24]. We reported earlier that these interactions are mediated by the intrinsically disordered regions (IDRs) present in these proteins [18].

Although a large body of evidence suggests a central role for the podocin molecule in the complex assembly, the precise mechanism by which podocin oligomerizes and acts as a scaffolding molecule remains to be elucidated. Importantly, it is not known whether if all the other SD proteins interact with a single podocin or to the homo-oligomers? Therefore, in this study, we attempted to understand the oligomeric nature of protein using a truncated construct (residues: 126-350), which encompasses the Prohibitin (PHB) domain, C-terminal oligomerization site, and 4 out of the 6 cysteines present in the native podocin sequence.

## Material and Methods

### Protein cloning, expression, and purification

The codon-optimized podocin gene (1152bp) was purchased from Gene Art (Life Technologies, USA). The regions encoding the amino acids 126-350 (podocin domain) was amplified with the primers 5’ CCC GAA TTC G AAA GTG GTG CAA GAA 3’ (forward) and 5’ GAA CTC GAG CAG ACA ATT CAG CAG ATC 3’ (reverse) and cloned into pET22b at EcoRI/XhoI sites. The recombinant construct was transformed into Arctic express (DE3) competent cells (Agilent Technologies, USA). The transformed cells were grown at 37°C in LB media supplemented with 100 μg/ml ampicillin and 10 μg/ml gentamycin. Protein expression was induced with 0.2 mM IPTG and cultured further for 16hrs at 14°C. The cells were harvested by centrifugation (13300 x g, 20 mins, and 4°C) and sonicated in the opening buffer (50 mM potassium phosphate (pH 8.0), 0.3 M NaCl, 5 mM β-mercaptoethanol and 0.1% Triton X-100) followed by clarification by centrifugation at 18000 x g for 45 mins at 4°C. The inclusion bodies were solubilized in 50 mM Potassium phosphate (pH 8.0), 0.3 M NaCl, 5 mM β-mercaptoethanol and 8 M Urea followed by clarification at 18000 x g for 1hr at room temperature. The solubilized protein was then purified using Ni-NTA agarose (Qiagen). The purity was confirmed (>98%) on a 12% SDS-PAGE. From the SDS-PAGE gel, the band corresponding to the podocin domain (27 kDa) was excised and subjected to trypsin digestion. The digested products were subjected to MALDI-TOF/TOF (Bruker Autoflex III smart beam, Bruker Daltonics, Bremen, Germany) to confirm the protein sequence.

The purified podocin was renatured by rapid dilution at 1:10 into 10mM Potassium phosphate buffer with 150 mM NaCl and 2 mM β-mercaptoethanol (pH 8.0) followed by dialysis in the same buffer to remove traces of urea and imidazole. The same pH and buffer composition are uniformly used in all the subsequent experiments. An excitation extinction coefficient of the 12950 M^−1^.cm^−1^ was used for determining the protein concentration on Jasco V-630 UV-Vis spectrophotometer

### Size exclusion chromatography and multi-angle light scattering analysis (SEC-MALS)

MALS in combination with SEC is a sensitive technique to accurately estimate the multiple oligomers within the sample when compared to SEC. We, therefore, performed SEC-MALS to estimate the oligomeric nature of the podocin domain. SEC-MALS was performed at room temperature by passing 500 μl protein (12μM) at 0.3 ml/min flow rate through a Superdex S200 SEC column (GE Healthcare) pre-equilibrated with 10mM potassium phosphate buffer supplemented with 150mM NaCl and 2mM β-mercaptoethanol (pH 8.0). This column was attached to the MALS system (AF2000-Postnova) for analyzing the molar mass of the protein. The protein sample from SEC-MALS was next passed through the flow cell equipped Zetasizer Nano ZS90 dynamic light scattering (DLS) device (Malvern Instruments Ltd, UK) equipped with a 4 mW He-Ne laser. The backscattering was measured at 173 nm for analyzing the polydispersity index (PDI) and the hydrodynamic radius of the protein.

### Structural features and Stability of the podocin domain

#### (a) Fluorescence spectroscopy

Intrinsic tryptophan fluorescence measurements were performed using a 1cm quartz cuvette on Jasco FP-6300 (Japan) equipped with an intense xenon flash lamp as the light source. 287 nm was used as the excitation wavelength and the 300-450 nm spectral range was used for obtaining the emission spectrum of the sample at 12 μM concentration. For accessing the stability of the podocin domain thermal-induced unfolding was performed. Fluorescence emission at 335 nm as a function of increasing temperature was recorded at a bandwidth of 2.5 nm and a scan speed of 200 nm/min with data recorded for every degree rise in temperature in triplicates. All the spectra were buffer corrected. The effect of temperature on the protein was analyzed by plotting fluorescence intensity at 335 nm using Origin (pro)-version2020b (Origin Lab Corporation, Northampton, MA).

#### (b) Circular dichroism spectroscopy

The circular dichroism (CD) measurements for podocin were recorded on Jasco J-1500 spectropolarimeter (Japan) equipped with a thermoelectric cell holder. The Far-UV (260-195 nm) CD measurements of the protein sample at 12 μM concentration were recorded using a 0.2 cm path length cell at 2.5 nm bandwidth and a scan speed of 50 nm/min. The Near-UV CD measurements were also recorded for the protein sample at 25 μM concentration using a 0.5cm pathlength cell at the bandwidth of 2.5nm and a scan speed of 100 nm/min. Both for far-UV and near-UV the data was accumulated in triplicates. To assess the effect of temperature and thus the stability of the domain, the sample was subjected to a steady increase in temperature and spectra were recorded at an interval of 5°C over a spectral range of 200 nm-250 nm. The data were plotted using origin lab software after buffer correction.

### Calorimetric analysis

Various thermodynamic properties including the melting temperature (Tm) of the podocin domain were obtained from measurements using NANO DSC (TA Instruments, USA). Sample containing protein concentration of 12 μM and a volume of 0.650 μL was loaded into the sample capillary and change in heat flow was recorded against reference buffer at a constant pressure of 3 atm, over a temperature range of 293K to 368K, with a scan rate of 1 K/min and a 300-sec cell equilibration time. Buffer scans were first performed before loading protein for baseline reproducibility. The obtained data was buffer corrected and the analysis data was plotted for peak integration with the peak analyzer option in Origin pro 2020b software. From the peaks, the transition temperature (T) and the enthalpy of unfolding (Δ_U_H) were calculated.

## Results

### Purification of Podocin domain

The region from 376bp to 1050bp of human *NPHS2* gene encoding 126-350 amino acid residues of podocin was PCR amplified and cloned into the pET22b expression vector (Fig. 1A&B). IPTG induced overexpression of the construct resulted in the protein to form inclusion bodies which were confirmed by Coomassie blue staining and immunoblotting with anti-His antibody (Fig. 2A&B). The protein was purified from inclusion bodies by Ni-NTA affinity chromatography after solubilization in 8M urea (Fig. 2C). Tryptic digestion of the band corresponding to the podocin domain from the SDS-PAGE gel (Fig. 2C-lane E2) and subsequent analysis of the digested products by MALDI-TOF/TOF revealed five peptide fragments (Fig. 2D and Table. 1). NCBI BLAST search of these peptide sequences against the non-redundant database confirmed the purified protein as human podocin covering the region 126-350 amino acids.

**Fig. 1:**
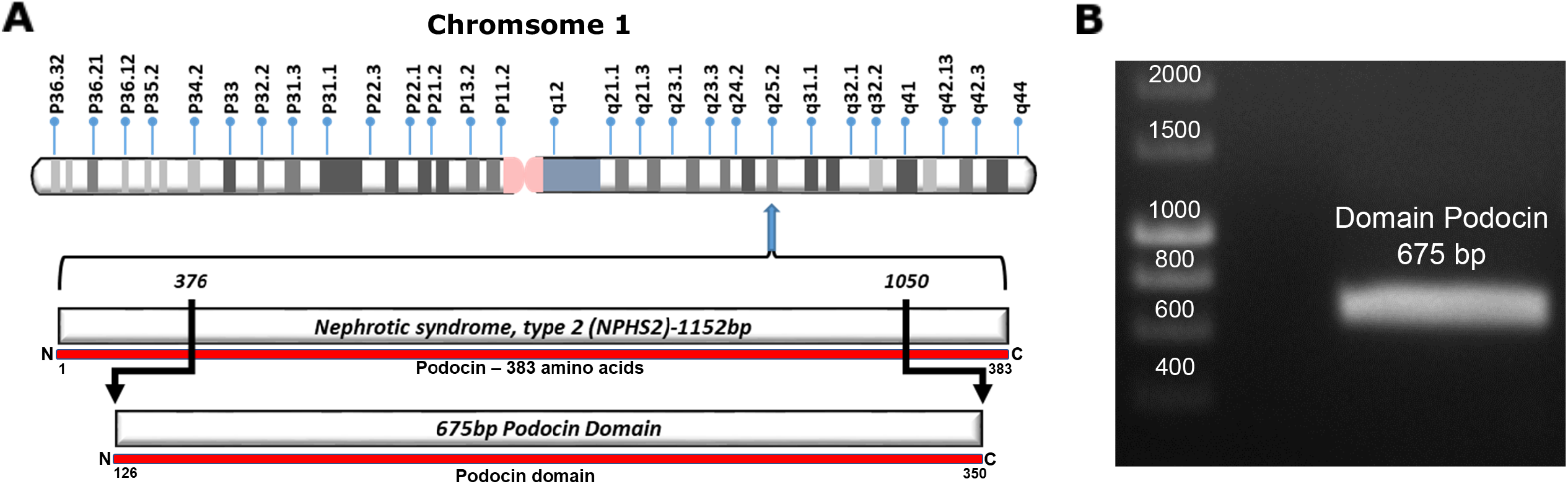
Cloning of podocin domain: A. The NPHS2 gene that encodes podocin is located on chromosome 1 at the locus q25.2. B. The region 376bp – 1050bp of the gene was amplified and cloned into pET22b bacterial expression vector.

**Fig. 2:**
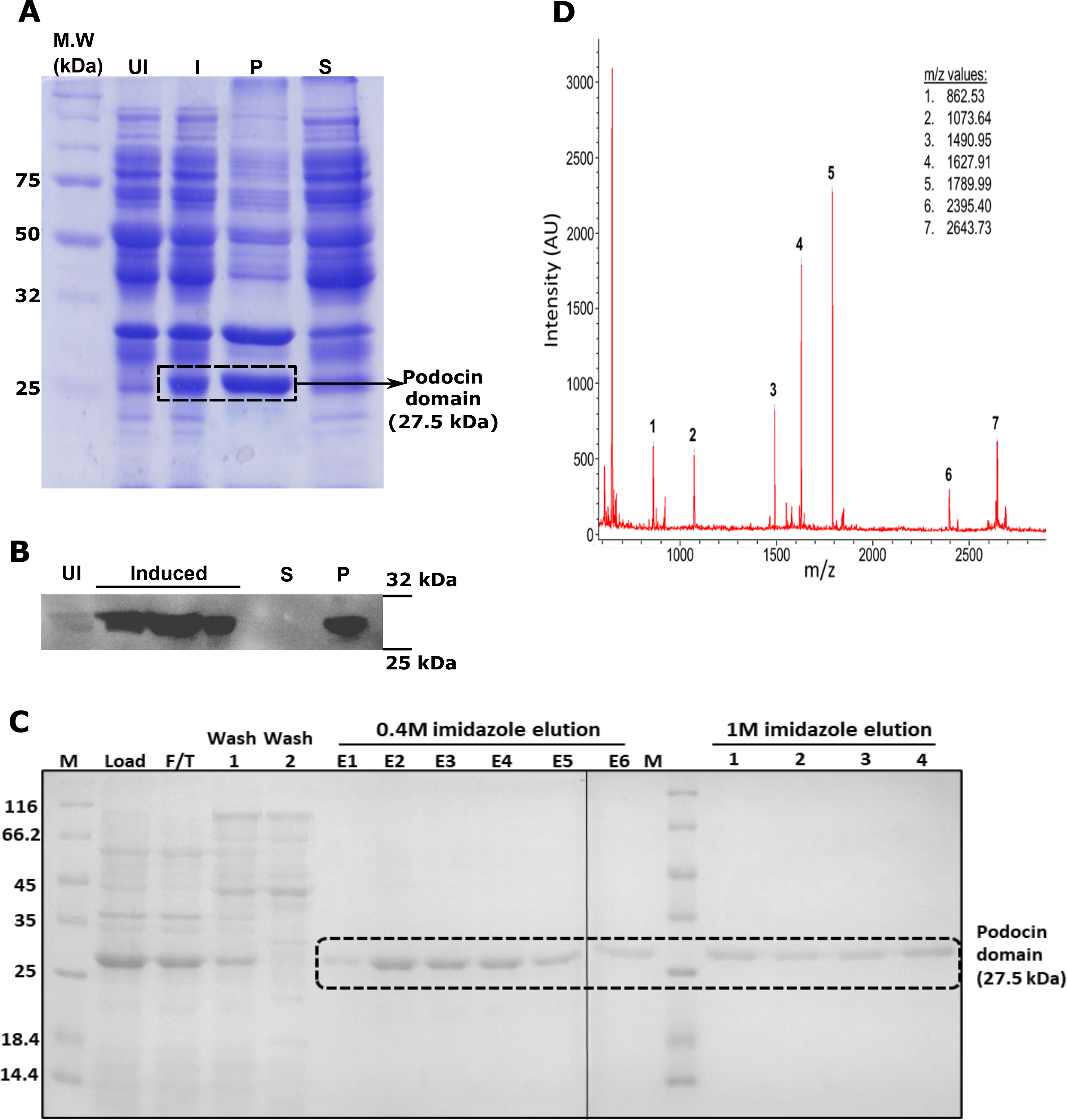
Expression, and purification of podocin domain: A & B. Coomassie blue staining and immunoblotting His-Tag HRP-conjugated antibody were done to identify the expression and solubility of the podocin domain. C. The SDS-PAGE analysis of samples after affinity chromatography purification of the urea solubilized cell lysate. D. Trypsinization and MALDI-TOF/TOF analysis of the 27KDa band excised from the earlier SDS-PAGE gel confirmed the presence of the podocin domain. UI-uninduced culture, I-Isopropyl β, D - thiogalactopyranoside Induced culture, P-Pellet fraction, S-Soluble fraction, F/T-flow through, E1-E6: elution fractions with 0.4M imidazole, 1-4: elution fractions with 1M imidazole.

**Table 1:**
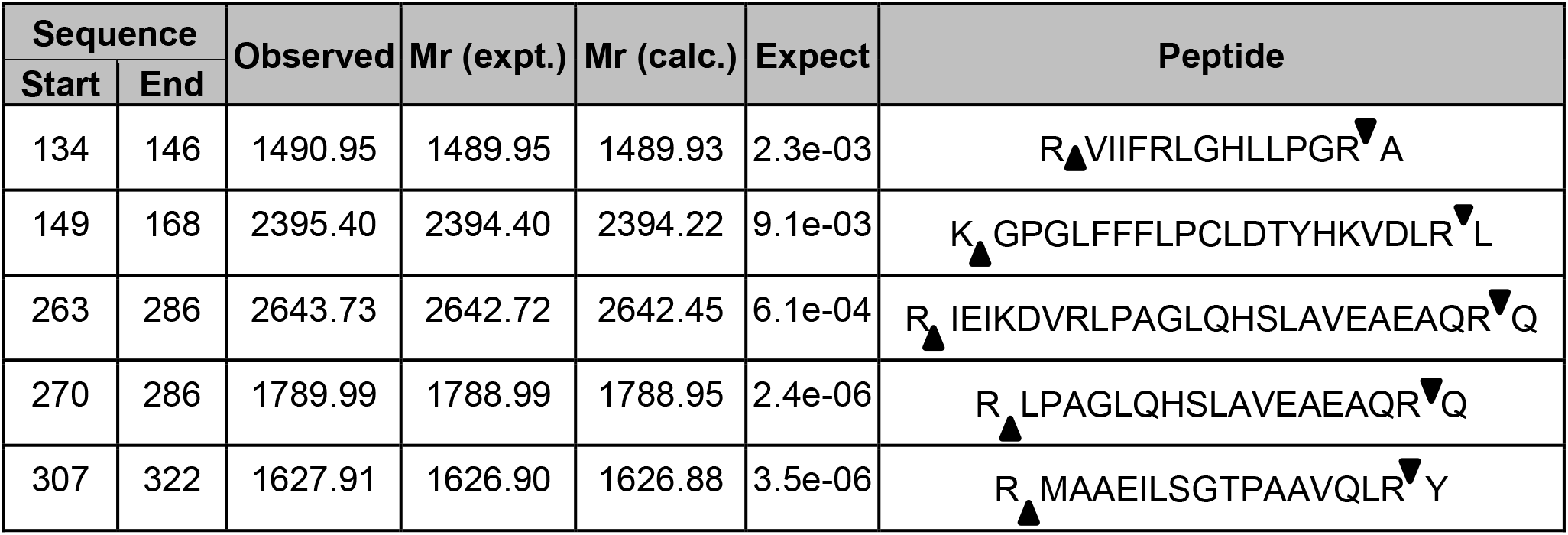
MALDI TOF/TOF analysis of the purified protein: Trypsinization of the purified band at 27 kDa and subsequence analysis by MALDI-TOF/TOF showed 5 peptide sequences. BLAST analysis of these sequences against the non-redundant proteins database of NCBI showed 100% similarity with the human podocin sequence. Note: ‘*▲*’ indicates the site of digestion by trypsin at arginine and lysine residues in the sequence.

### Oligomeric nature of podocin domain

It was reported that stomatin family members exist as homo-oligomers [20–22]. Since podocin shares significant homology with stomatin, we analyzed the oligomeric nature of the podocin domain consisting of the PHB domain. SEC-MALS data is represented as a combinatorial plot of refractive index, and molecular weight vs elution time (Fig. 3). A maximum refractive index value of 0.11 for the podocin domain was observed which corresponds to a molecular weight of 450kDa, suggesting that the podocin domain is a 16mer oligomer (Molecular weight of monomer = 27.5kDa; therefore; 450 kDa/ 27.5 kDa = ~16mer). In addition to the predominant 16mer species, other oligomeric conformations of the podocin domain ranging from 25mer (refractive index: 0.08, molecular weight: 697kDa) to 13mer (refractive index: 0.10, molecular weight: 376kDa) were also observed, but to a lesser extent. The DLS, which was in tandem with the SEC-MALS analyzed the eluted samples for hydrodynamic radius and polydispersity. A hydrodynamic radius range of 13.39 - 9.37 nm corresponding to elusions from 25mer to 13mer was observed respectively (Fig. 4A-E). The average PDI of the sample was found to be 0.14, which suggests that the sample consists of one major population by volume, however, a broad size range within the population implies the presence of different size species. From these results, it is evident that the major species the podocin domains associate into 16mer oligomers while a minor population of other higher-order oligomers also appear to exist in solution.

**Fig. 3:**
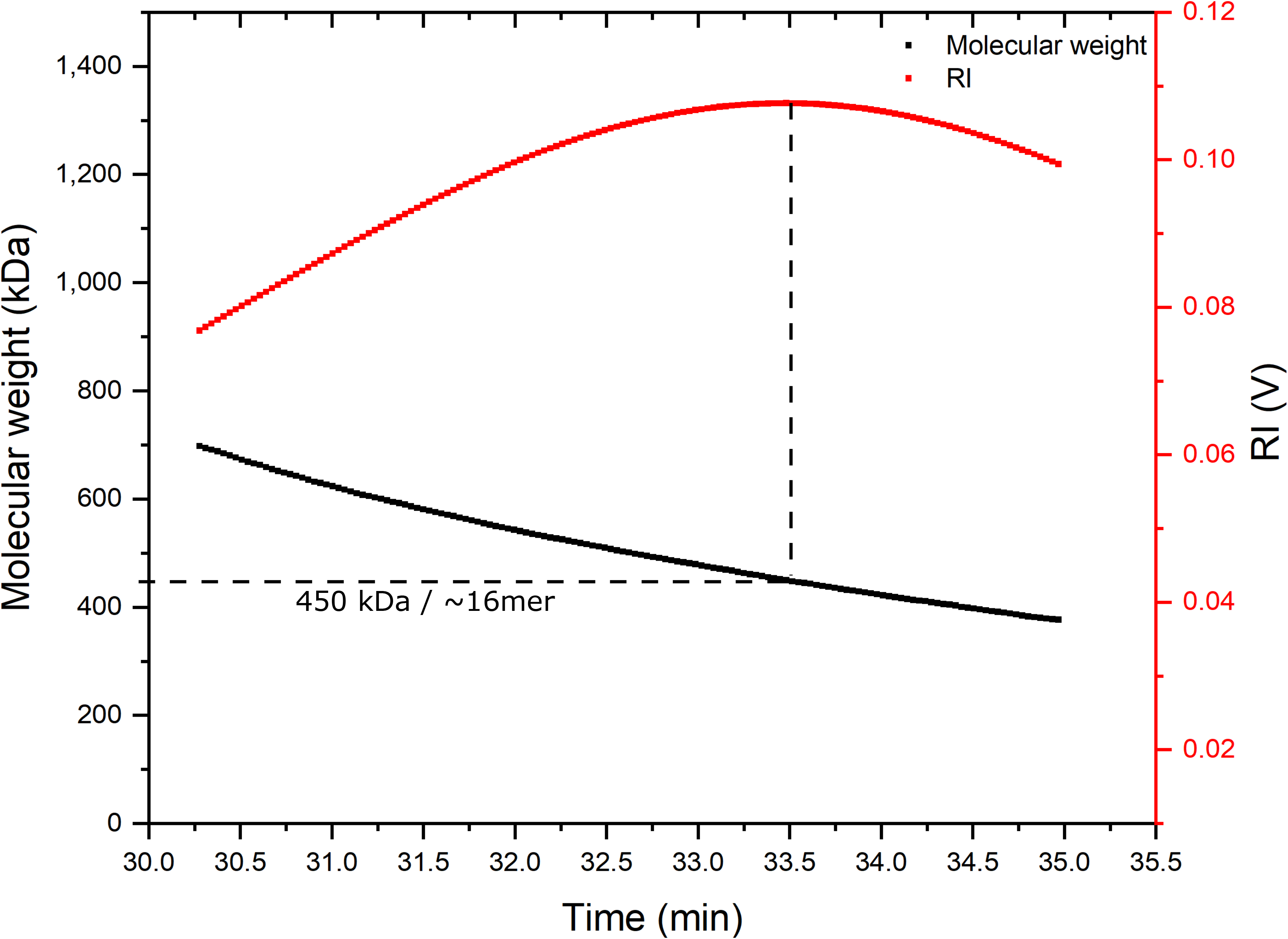
Podocin domain forms higher-order oligomers: SEC-MALS analysis of the podocin domain for molecular mass determination. The molecular weight on Y-axis and the refractive index on the secondary axis and were plotted against elution time.

**Fig. 4:**
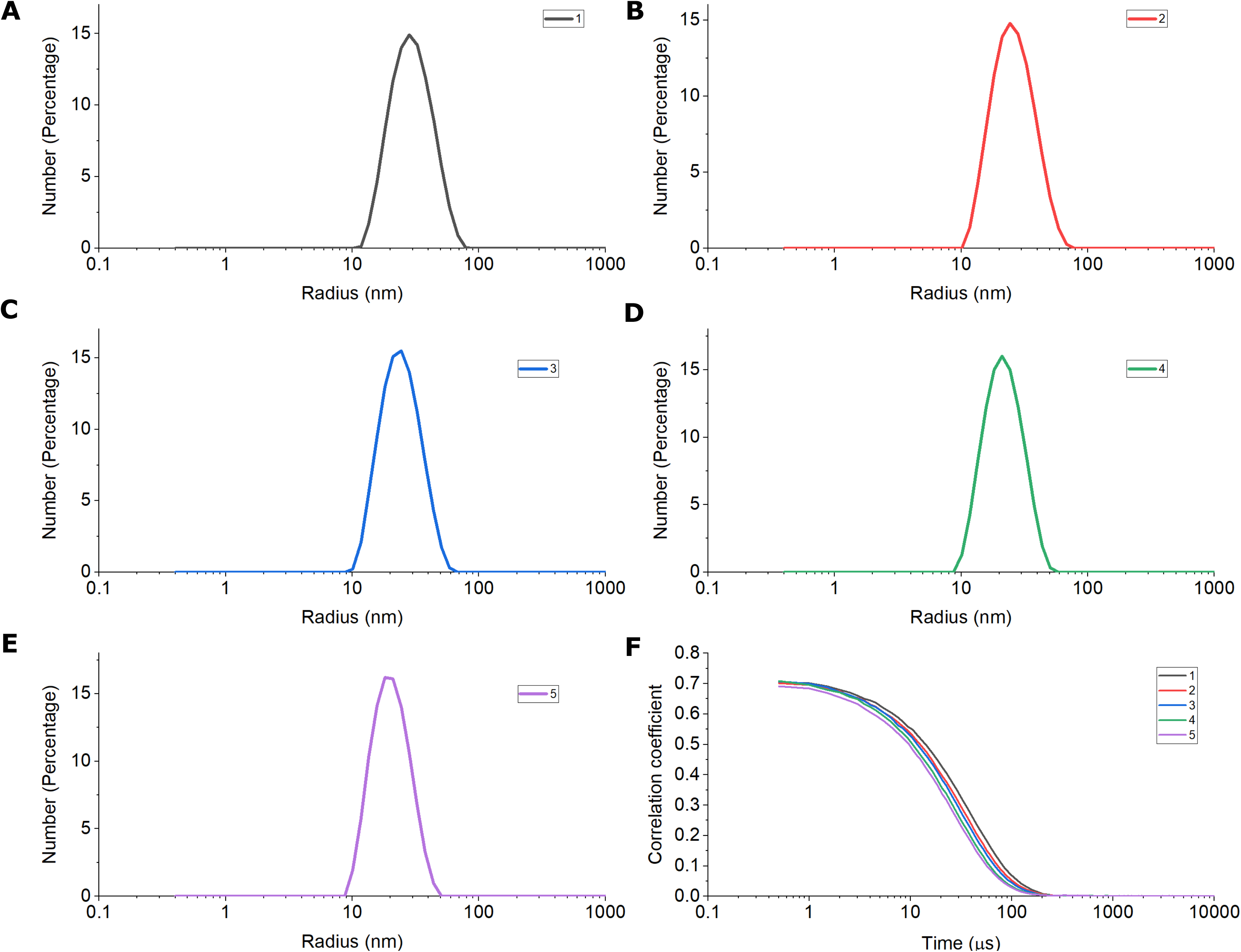
Polydispersity and hydrodynamic radius of the podocin domain homo-oligomers: The DLS was in tandem with the SEC-MALS. DLS analysis of the samples corresponding to peak observed in SEC-MALS (1-5) is represented as a number percentage vs size in nanometers curve plots marked in different colors (A-E). The corresponding correlograms of the samples (1-5) represented as correlation coefficient vs time in microseconds.

### Structural features of the homo-oligomers of podocin domain

CD spectra at ambient temperature in the far (250–195 nm) and near-UV region (310–250 nm) were recorded (Fig. 5A&B). The far UV region provides information on the secondary structure while the near UV region on the tertiary interactions. Observation of CD signal in the near UV regions implies the arrangement of aromatic amino acids in a restricted environment(s) and thus implying a folded structure adopted by the polypeptide chain. The far UV CD signal for the truncated podocin domain indicates that the protein adopts secondary structure. The shape of the spectrum indicates the possibility of the presence of both α helices and β sheet structures. The near UV spectrum of the sample shows a broad peak at 280nm typically contributed from tryptophan residue (W256), and a minor peak from ~250 – 260 nm could be from five-tyrosine and eight-phenylalanine residues. The presence of signals in this region implies the tertiary interactions involving these residues.

**Fig. 5:**
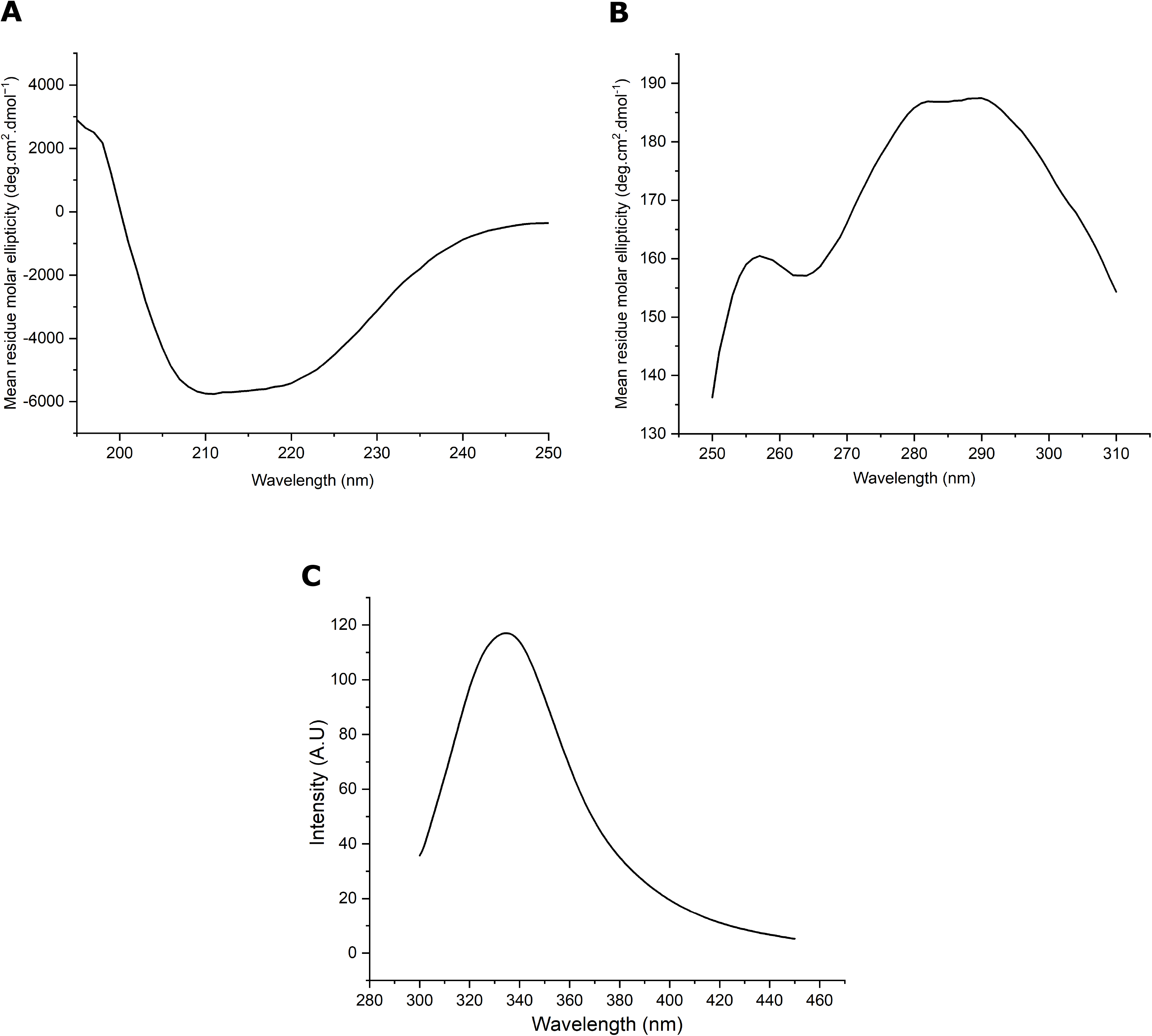
Podocin domain exhibits secondary structural elements and tertiary packing: A. Far-UV CD spectrum of Podocin domain at 25°C, pH 8 in 10mM potassium phosphate buffer supplemented with 150mM NaCl and 2mM β-mercaptoethanol. B. Near-UV CD spectrum representing the tertiary packing of the podocin domain. C. Fluorescence emission of the podocin domain recorded by exciting the protein at 287 nm.

The intrinsic tryptophan emission spectrum of the podocin domain shows a λ_max_ value at 335nm (Fig. 5C). The λ_max_ of native proteins relates to the polarity of the environment of the tryptophan residue and typically can range from 308-350 nm. Unfolded forms with residues in the apolar microenvironment show a blue shift of the λ_max_ [25]. Although, the intrinsic emission may not directly or unequivocally provide structural information, changes in the folded state can be followed by monitoring the changes in the emission intensity which convey the changes in the native state tryptophan environment. The changes in the CD signal and the perturbations of the intensity of the intrinsic fluorescence emission were used to estimate the stability of the folded podocin domain.

### Structural stability of the podocin domain

To assess the structural stability, the resistance of the structure to unfold on heating was monitored following changes in the tryptophan fluorescence emission and the far UV CD spectrum. Temperature-induced unfolding was monitored using the fluorescence emission spectra with increasing temperature in the range of 20-95°C. The spectra showed a uniform decrease in λ_max_ intensity without either a bathochromic or hypsochromic shift (Fig. 6A). The change in fluorescence intensity as a function of temperature (Fig. 6B) shows a linear transition and does not show a sigmoidal shape typically observed for proteins with a tightly packed tertiary core. The absence of a folded baseline suggests a not so tightly packed tertiary core. Far UV CD spectral changes were also monitored as a function of temperature. The signal intensity showed a gradual loss with increasing temperature (Fig. 6D), and the thermal unfolding monitoring the loss of secondary structure also showed the absence of a native baseline (Fig. 6C) implying that the podocin domain possesses secondary and tertiary interactions but lack a tightly packed core.

**Fig. 6:**
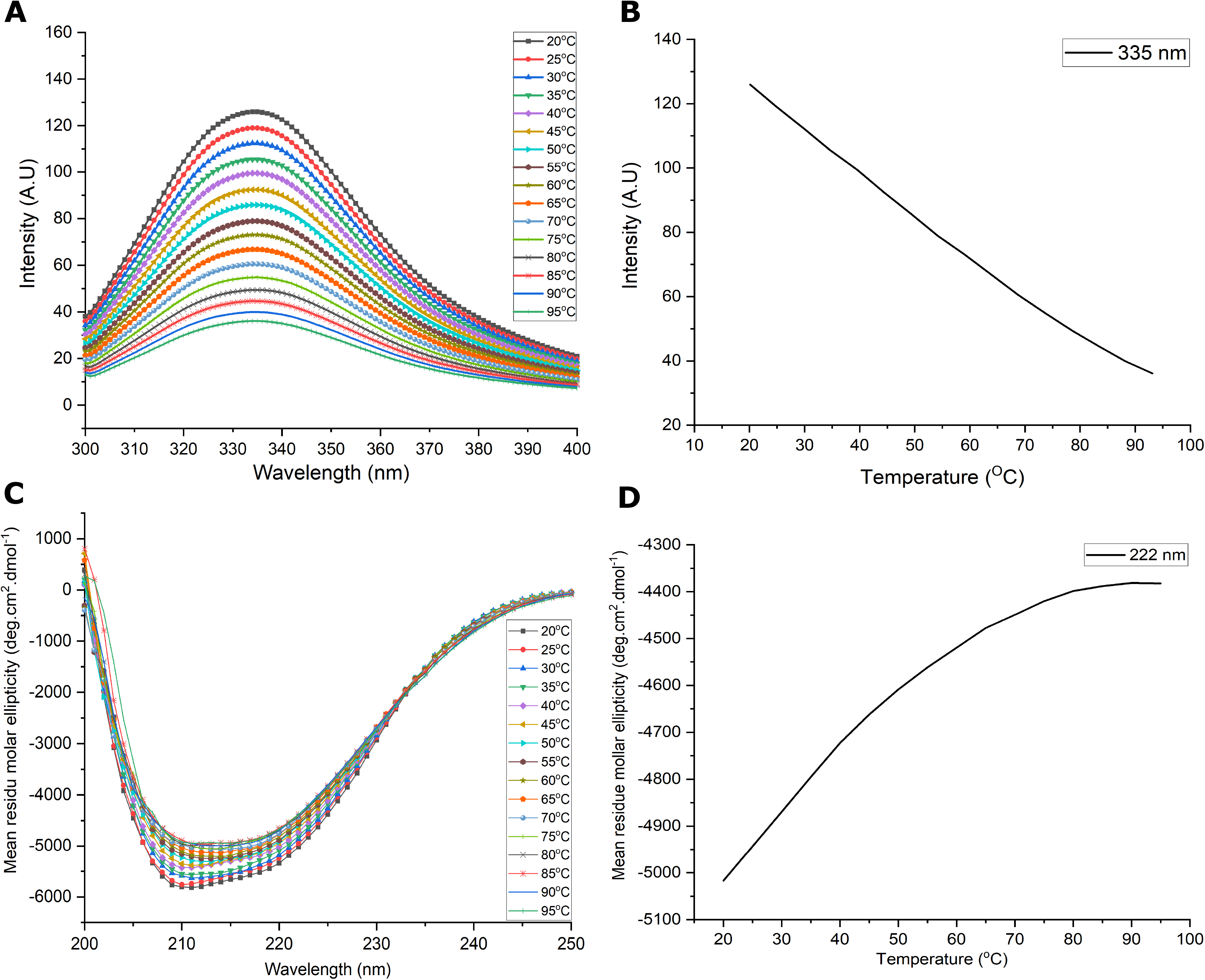
Effect of temperature on podocin domain: A. Intrinsic tryptophan fluorescence for podocin domain was measured as a function of temperature (20°C to 95°C) by exciting the protein at 287 nm. B. Peak emission of the podocin domain at 332 nm is plotted over a temperature range of 20°C to 95°C. C. FarUV CD spectra (250 – 200 nm) of the podocin domain was acquired as a function of temperature (20°C to 95°C). D. The MRE values of the podocin domain at 222nm is plotted as a function of temperature (range: 20°C to 95°C) to monitor the changes in the protein structure.

### Calorimetric analysis of podocin domain

Differential scanning calorimetry (DSC) gives the overall enthalpy value (ΔH_cal_) for each structural transition. Therefore, we performed DSC to calculate the thermodynamic parameters such as transition temperature (T) and the enthalpy of transition associated with structural changes of the homo-oligomer. Peak deconvolution of the acquired data revealed 5 transition states, out of which four are endothermic transitions (316 K, 325 K, 330 K, and 353 K) and one is an exothermic transition (345 K) (Fig. 7). The respective values for ΔH are mentioned in Table 2. The DSC profile suggests that the oligomers in the mixture undergo dissociation via three transition temperatures namely 316 K, 325 K, and 330 K and the presence of exothermic transition at 345 K suggests possible hydrophobic interactions among the constituent oligomers before complete dissociation at 353 K.

**Fig. 7:**
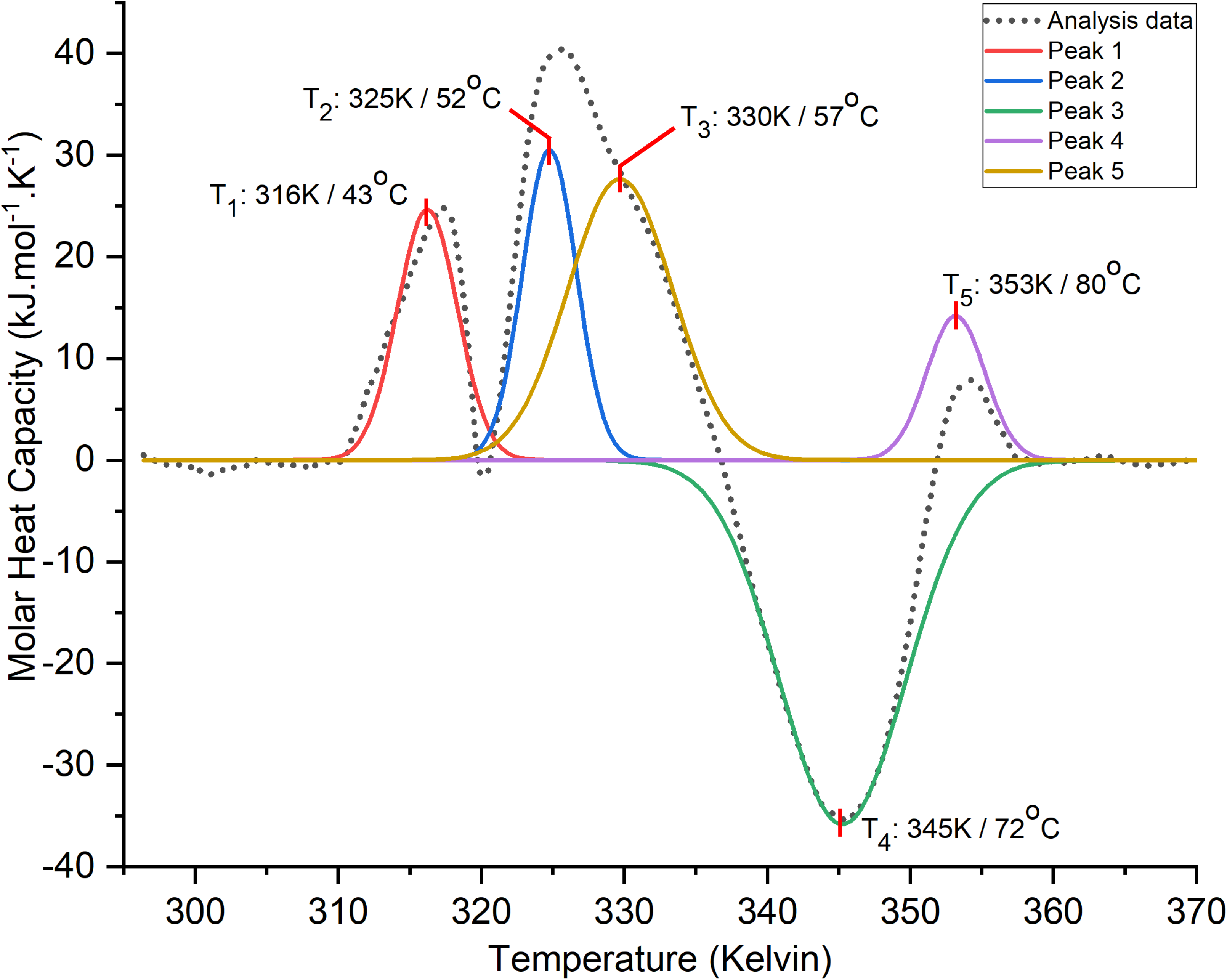
DSC analysis of podocin domain: The plots represent the endothermic and exothermic transitions of the podocin domain over a temperature range from 296K/23°C to 368K/95°C. The initial DSC artifact near 293K/20°C in the DSC data is not represented in the plot. The analysis data represented as a dotted line in the plot after baseline subtraction. Each transition peak (1-5) in the plot is represented by a color line which is a result of peak deconvolution function.

**Table. 2:** Enthalpy values and the transition temperatures as noticed in the dynamic scanning calorimetry.

| Temperature range | Transition temperatures | | Enthalpy change ($\Delta H_{cal}$) (kJ/mol) |
| --- | --- | --- | --- |
| 293K to 368K /<br>(20°C – 95°C) | Endothermic | 316 K / 43°C | 130 |
|  |  | 325 K / 52°C | 145 |
|  |  | 330 K / 57°C | 73 |
|  |  | 353 K / 80°C | 254 |
|  | Exothermic | 345 K / 72 °C | -397 |

## Discussion

Podocin selectively expresses in the glomerular podocytes and it is instrumental for preserving the structural integrity of the SD. Though several mutations in the protein are associated with proteinuria in humans, the structural details of this protein are unclear. Here we report for the first time, the stoichiometry of oligomerization of the truncated human podocin construct. Our investigation indicates that at ambient temperature, and in a reduced environment the podocin domain predominantly adopts a 16mer oligomeric state. However, other oligomeric conformations ranging from 25mer to 13mer were also observed nevertheless, the population of these states was comparatively less. The polydispersity index we report adds evidence to the presence of multiple oligomeric species. Additionally, through CD and Fluorescence spectrums we show that the homo-oligomers have considerable structure and tertiary interactions.

Since stomatin proteins and podocin share homology, it expected they may share several structural similarities. Crystallization studies of stomatin protein (*Pyrococcus horikoshii*; residues: 56–224) revealed that it exists as a trimer and NMR studies of the same protein but with a different truncation size (residues: 66–174) associated into an amalgam of oligomers [20]. Earlier reports have suggested that podocin forms homo-oligomers with its C-and N-terminals at the lipid raft micro-domains besides forming a large complex with its neighboring SD proteins [7, 16, 17, 26]. However, the extent of the oligomer size was not explored. Furthermore, a recent study by Straner et.al, reported that shorter C-terminal construct of human podocin containing the PHB domain formed homodimers via helical segments (283–313 and 332–348 residue. Additionally, they have also noted that the full C-terminal construct (168–383) formed higher-order oligomers [23]. However, insights into the mechanism of oligomeric assembly and stoichiometry remain largely unknown.

We were unsuccessful in expressing and purifying the full-length human podocin protein. This could be due to several reasons including podocin may not be stable outside its native environment, due to the presence of IDRs, or interference of the transmembrane segment with protein expression [5]. We, therefore, cloned and expressed a truncated construct encoding the 126-350 amino acids region of the podocin. This region encompasses the PHB domain and the C-terminal oligomerization site. SEC-MALS analysis of the protein revealed that the podocin domain associated majorly into 16mers. It also adopted other oligomeric states ranging from 25-13mers. These results are further justified by the hydrodynamic radius and the polydispersity index measured by the DLS. The PDI reported for the podocin domain falls into the mid-range i.e., a value between 0.08 – 0.5 suggests that the protein sample although has one major oligomer species by volume, the presence of multiple species is possible [27]. Due to the presence of multiple oligomeric species and apparent dynamicity between the molecules of the oligomer, we were unable to obtain crystals of the podocin domain for performing X-ray diffraction. Nevertheless, our study adds evidence to the observations made in earlier reports that longer C-terminal constructs of podocin may adopt higher oligomeric conformations [17, 23].

Far UV-CD spectra revealed that the podocin domain consists of a considerable amount of α-helices and β-sheets. We could not record far UV-CD spectra beyond 195nm possibly because of the presence of buffer components like β-mercaptoethanol that hindered the spectrum which is reflected as increased HT voltage. The inclusion of β-mercaptoethanol in the buffer helped in solubilizing podocin domain and contributed towards the protein stability. It is noteworthy that crystal structures of stomatin protein are devoid of disulfide linkages [22, 28]. In a study, Huber et.al reported that Cys126 and Cys160 residues of mouse podocin undergo palmitoylation and participate in membrane insertion [26]. Similarly, the Cys158 in the human podocin domain is also expected to be palmitoylated instead of forming disulfide linkages. This suggests that the disulfide bridges between cysteines either do not exist or are not contributing to protein stability.

While assessing the effect of temperature on the protein by probing its intrinsic tryptophan fluorescence and secondary structure content via measuring optical rotation by using the CD, the baseline was observed only in the case of CD spectrum plots beyond 80°C. A significant linear decrease in the fluorescence intensity at 335 nm with increasing temperature may be due to the partial exposure of the lone Trp256 and other aromatic residues. Also, there was no bathochromic shift observed which ascertains the fact that the oligomer might not have dissociated completely. Similarly, far UV – CD spectra at 222nm was used to monitor secondary structure changes. A small yet noticeable change in the MRE value and the shape of the spectra implicates that most of the secondary structure was retained, however, a slight rearrangement of in structure could be possible, owing to the increased hydrophobic effect. With the increasing temperature, the prominent double minima smoothen out indicating that there might be a slight increase in the beta-sheet content. The stable MRE value observed beyond 80°C can be the mark for the stabilization of some conformer.

Calorimetric analysis of the oligomeric species revealed multiple transition points for the podocin domain. At 20°C the protein was shown to attain an amalgam of oligomeric states and a steady increase in temperature may promote dissociation of these different oligomers via three different transition temperatures since each oligomeric state may not have same transition point. Nevertheless, when the temperature was further increased significant exothermic transition state was observed. We assume the exothermic transition state may be due to the association of hydrophobic cores of the constituent oligomers. It is well known that lower temperatures do not favor hydrophobic interactions whereas, an increase in temperature to a certain degree promotes hydrophobic interactions among the protein molecules [29]. Subsequently, when the temperature is further increased, we speculate that the oligomer may invariably disassociate into lower oligomeric species. It is noteworthy that the podocin domain did not re-trace the path to its initial state when the protein was cooled. This peculiar behavior of the podocin domain starkly correlates with the oligomeric propensities observed in the stomatin proteins [20, 21].

All the above-discussed experiments were also performed at 5°C to ensure whether if podocin domain attains a stable 16meric oligomeric state as a predominant conformation in its energy landscape. From the data, we could not observe much difference in terms of the major species populating in the protein sample. 16mer seems to be the major/stable conformer at both the temperatures (Fig S1-S4). Interestingly in the calorimetric analysis, multiple transition points were not observed which might confirm the presence of 16mer oligomeric species as the predominant conformer at lower temperatures which was further confirmed by SEC analysis (Fig. S3-S4).

In conclusion, our study, for the best we know is the first to report the cloning, expression, and purification of the human podocin domain (126-350). We showed that the podocin domain is majorly a 16mer homo-oligomer. However, to a lesser extent, the protein is capable of associating into other oligomeric states. CD and FL analysis of the protein indicated that the podocin domain in isolation attains considerable secondary structure and tertiary packing. However, the significance of the podocin oligomerization in the formation of the large macromolecular complex and how a mutation in a podocin monomer will affect its oligomerization status need to be addressed. Further structural characterization of podocin and other slit diaphragm proteins is greatly warranted to understand the mechanism of pathogenesis of the nephrotic syndrome.

## Acknowledgments

We acknowledge Dr. Akif, University of Hyderabad, for providing access to the AKTA-Purifier Gel-filtration system. We also acknowledge the Central Analytical laboratory of BITS - Pilani, Hyderabad campus for allowing access to Circular Dichroism Spectrophotometer. MNSK acknowledges ICMR for providing senior research fellowship.

## Author Contributions

MNSK, AKP, KG, VP, and RV planned experiments; MNSK, KK, and SSI performed experiments; MNSK, AKP, KG, VP, and RV analyzed data; AKP, VP, and RV contributed reagents; MNSK, AKP, SSI, and RKV wrote the paper.

## Abbreviations

SD: slit-diaphragm
GFB: Glomerular filtration barrier
CD2AP: CD-2 associated protein
TRPC6: Transient receptor potential cation channel subfamily C member 6
ZO-1: Zonula occludens-1
NEPH: Nephrin-like protein
NS: Nephrotic syndrome
SRNS: steroid-resistant NS
*NPHS1* & 2: Nephrotic syndrome-type I and type II
IDRs: Intrinsically disordered regions
SEC: Size-exclusion chromatography
MALS: multi-angle light scattering
CD: Circular dichroism

## Supplementary Figures

**Fig. S1: Podocin domain exhibits secondary structural elements:** Far-UV CD spectrum of Podocin domain at 5°C, pH 8 in 10mM potassium phosphate buffer supplemented with 150mM NaCl and 2mM β-mercaptoethanol.

**Fig. S2: Effect of temperature on podocin domain:** A. Intrinsic tryptophan fluorescence for podocin domain was measured as a function of temperature (5°C to 95°C) by exciting the protein at 287 nm. B. Peak emission of the podocin domain at 332 nm is plotted over a temperature range of 5°C to 95°C. C. FarUV CD spectra (250 – 200 nm) of the podocin domain was acquired as a function of temperature (5°C to 95°C). D. The MRE values of the podocin domain at 222nm is plotted as a function of temperature (range: 5°C to 95°) to monitor the changes in the protein structure.

**Fig. S3**: **Podocin domain majorly adopts 16mer homo-oligomer**: SEC of the purified podocin domain at 4°C showing different podocin domain oligomeric conformations. A pre-packed HiLoad 16/600 Superdex 200pg (GE Healthcare) column was equilibrated with 10 mM potassium phosphate, 150 mM NaCl, and 2 mM β-mercaptoethanol (pH 8.0). 1ml of protein at 9 μM concentration was applied to the column at a constant flow rate of 0.5ml/min. The elution volume of the protein was detected by monitoring the UV absorbance at 280 nm.

**Fig S4: DSC analysis of podocin domain:** The plots represent the endothermic and exothermic transitions of podocin domain over a temperature range from 5°C to 95°C. The initial DSC artifact near 273K in the DSC data is not represented in the plot. The analysis data after baseline subtraction is represented as a dotted line in the plots, whereas the transition peaks in the data are represented as solid color lines.

