## Supplementary figures and images for "Structural Features and Oligomeric Nature of Human Podocin Domain"

### Fig S1-S4

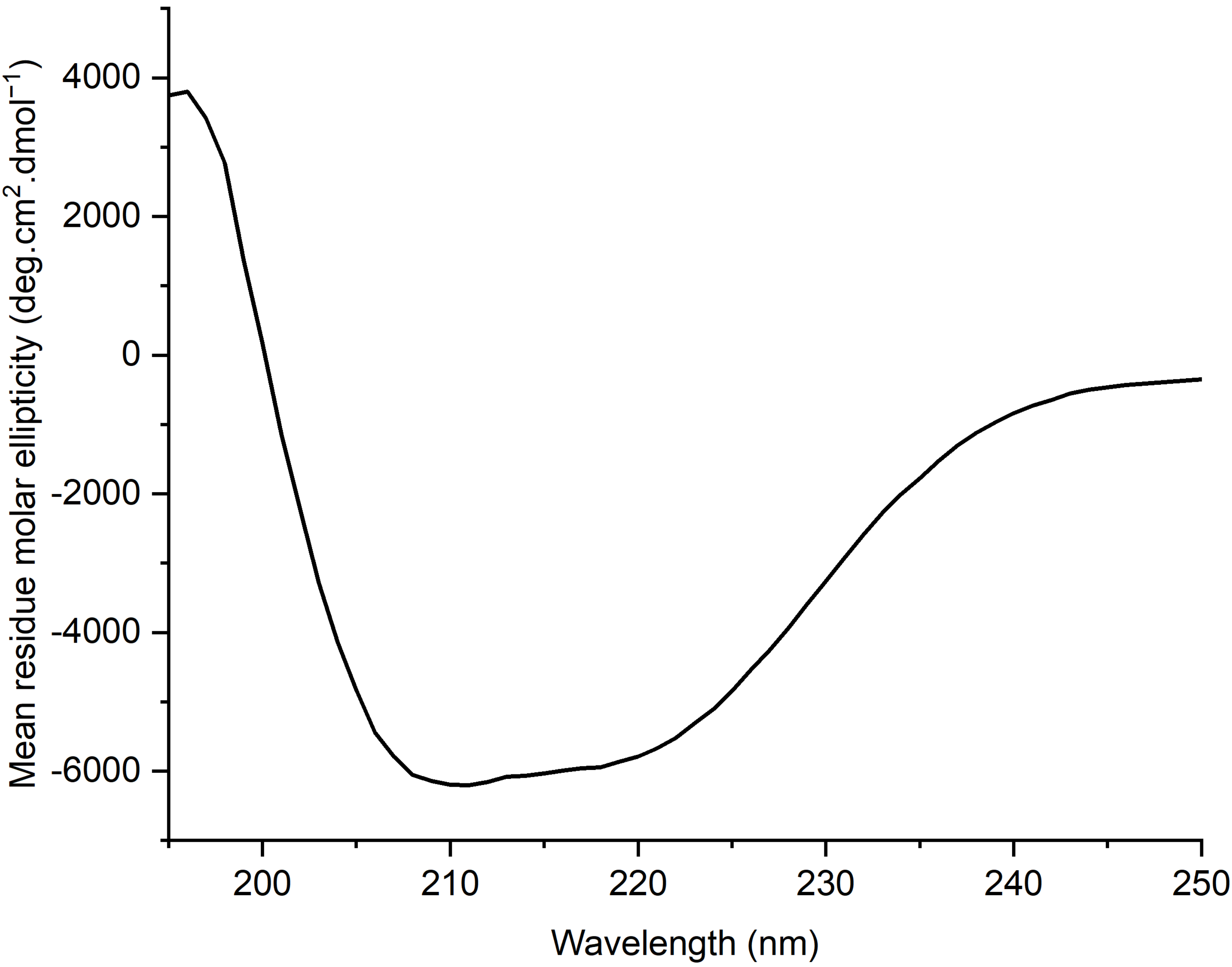

### Fig S1-S4

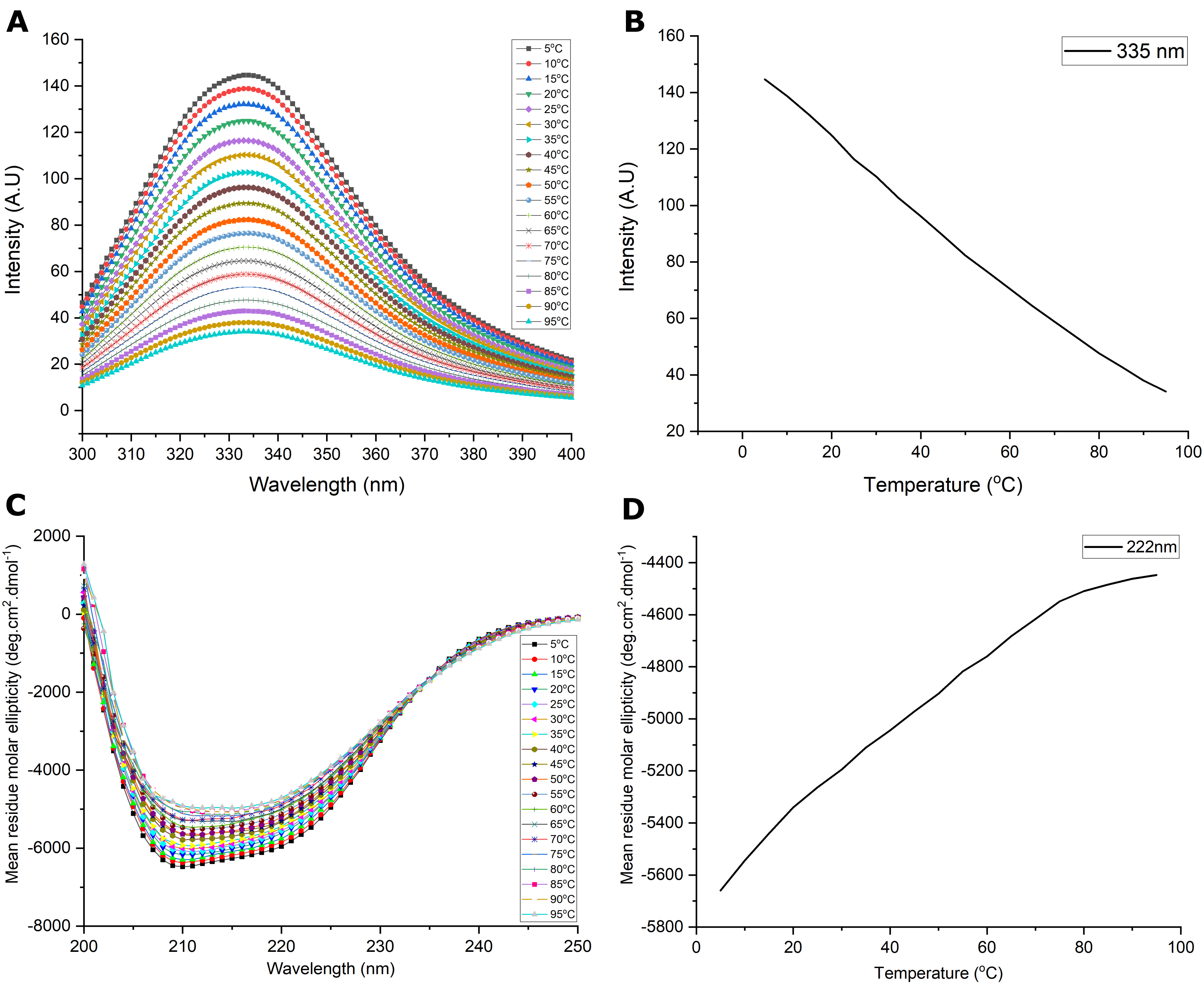

### Fig S1-S4

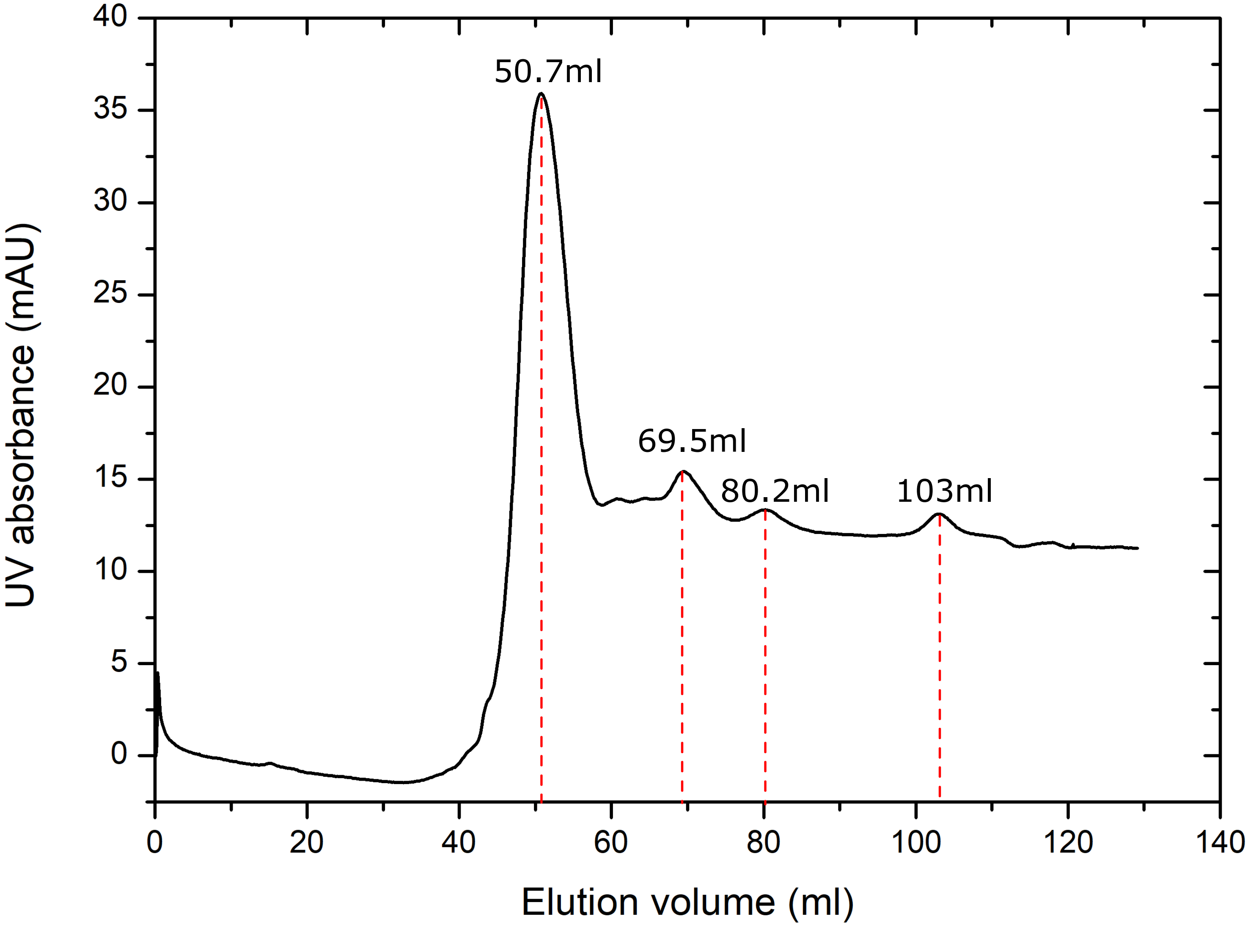

### Fig S1-S4

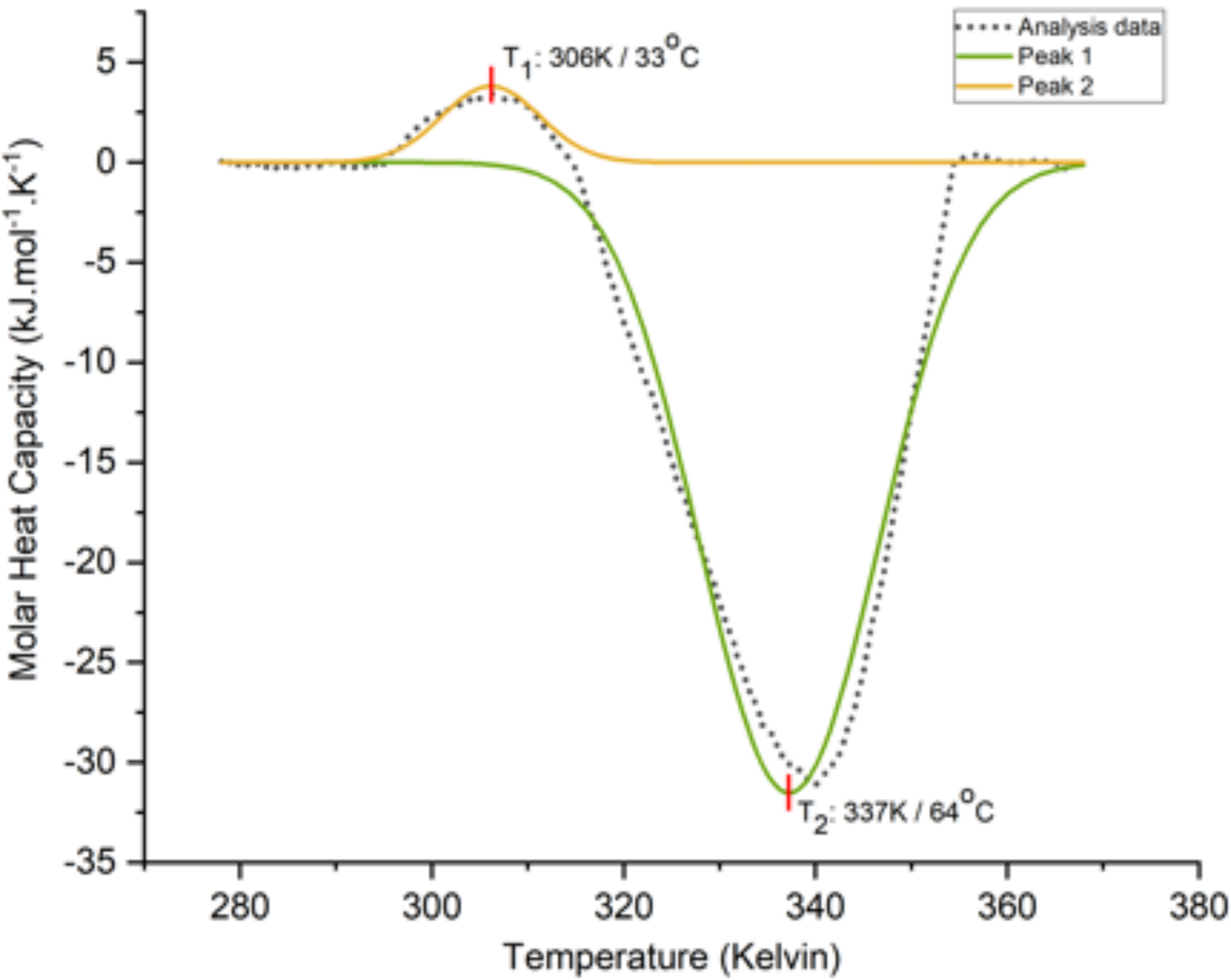
